# Bivalent SARS-CoV-2 mRNA vaccines increase breadth of neutralization and protect against the BA.5 Omicron variant

**DOI:** 10.1101/2022.09.12.507614

**Authors:** Suzanne M. Scheaffer, Diana Lee, Bradley Whitener, Baoling Ying, Kai Wu, Hardik Jani, Philippa Martin, Nicholas J. Amato, Laura E. Avena, Daniela Montes Berrueta, Stephen D. Schmidt, Sijy O’Dell, Arshan Nasir, Gwo-Yu Chuang, Guillaume Stewart-Jones, Richard A. Koup, Nicole A. Doria-Rose, Andrea Carfi, Sayda M. Elbashir, Larissa B. Thackray, Darin K. Edwards, Michael S. Diamond

**Affiliations:** Department of Medicine, Washington University School of Medicine, St. Louis, MO 63110, USA; Moderna, Inc., Cambridge MA, USA; Vaccine Research Center, National Institute of Allergy and Infectious Diseases, Bethesda, MD, USA; Department of Pathology & Immunology, Washington University School of Medicine, St. Louis, MO, USA; Department of Molecular Microbiology, Washington University School of Medicine, St. Louis, MO, USA; The Andrew M. and Jane M. Bursky Center for Human Immunology and Immunotherapy Programs, Washington University School of Medicine. St. Louis, MO, USA; Center for Vaccines and Immunity to Microbial Pathogens, Washington University School of Medicine, Saint Louis, MO, USA

## Abstract

The emergence of SARS-CoV-2 variants in the Omicron lineage with large numbers of substitutions in the spike protein that can evade antibody neutralization has resulted in diminished vaccine efficacy and persistent transmission. One strategy to broaden vaccine-induced immunity is to administer bivalent vaccines that encode for spike proteins from both historical and newly-emerged variant strains. Here, we evaluated the immunogenicity and protective efficacy of two bivalent vaccines that recently were authorized for use in Europe and the United States and contain two mRNAs encoding Wuhan-1 and either BA.1 (mRNA-1273.214) or BA.4/5 (mRNA-1273.222) spike proteins. As a primary immunization series in BALB/c mice, both bivalent vaccines induced broader neutralizing antibody responses than the constituent monovalent vaccines (mRNA-1273 [Wuhan-1], mRNA-1273.529 [BA.1], and mRNA-1273-045 [BA.4/5]). When administered to K18-hACE2 transgenic mice as a booster at 7 months after the primary vaccination series with mRNA-1273, the bivalent vaccines induced greater breadth and magnitude of neutralizing antibodies compared to an mRNA-1273 booster. Moreover, the response in bivalent vaccine-boosted mice was associated with increased protection against BA.5 infection and inflammation in the lung. Thus, boosting with bivalent Omicron-based mRNA-1273.214 or mRNA-1273.222 vaccines enhances immunogenicity and protection against currently circulating SARS-CoV-2 strains.

## INTRODUCTION

The SARS-CoV-2 pandemic has caused more than 600 million infections and 6.4 million deaths (https://covid19.who.int). In response to the global public health challenge, multiple companies rapidly developed vaccines using several different platforms *(e.g*., (lipid nanoparticle encapsulated mRNA, inactivated virion, nanoparticle, or viral-vectored vaccine); some of these vaccines have been approved by regulatory agencies in different parts of the world and deployed in billions of people, resulting in reduced numbers of infections, hospitalizations, and COVID-19-related deaths. The target antigen for most of these SARS-CoV-2 vaccines is the spike protein derived from historical strains that circulated in early 2020. However, the continuing evolution of SARS-CoV-2, resulting in amino acid changes in the spike protein amidst successive waves of infection, has jeopardized the efficacy of global vaccination campaigns and the control of virus transmission^1^.

The SARS-CoV-2 spike protein binds to angiotensin-converting enzyme 2 (ACE2) on human cells to facilitate viral entry and infection^2^. The S1 fragment of the spike protein contains the receptor binding domain (RBD), which is the primary target of neutralizing antibodies elicited by vaccination or produced after natural infection^3-5^. In late 2021, the first Omicron variants (BA.1 and BA.1.1) emerged, with greater than 30 amino acid substitutions, deletions, or insertions in the spike protein. Since then, the Omicron lineage has continued to evolve (*i.e*., BA.2, BA.4, BA.5, BA.2.75, and BA.4.6) with additional or different sets of spike mutations that facilitate escape from neutralizing antibodies^6,7^. These changes in the spike protein of Omicron strains are associated with symptomatic breakthrough infections in vaccinated and/or previously infected individuals^8-10^.

To overcome the loss in efficacy of the vaccines against Omicron strains, third and even fourth doses (herein, boosters) of mRNA vaccines encoding the historical (Wuhan-1) spike protein were recommended, and vaccines with Omicron variant-matched spikes were rapidly designed and tested. In humans, a booster dose of mRNA-1273 vaccine was associated with neutralizing antibody titers against BA.1 that were approximately 20-fold higher than those assessed after the second dose of vaccine^11^. In both mice and non-human primates, boosting with either mRNA-1273 or an Omicron BA.1-matched (mRNA-1273.529) vaccine increased neutralizing titers and protection against BA.1 infection compared to animals given a primary (two-dose) vaccination series of mRNA-1273^12,13^. Moreover, neutralizing antibody titers were higher, and BA.1 viral burden in the lung was lower, in mice boosted with mRNA-1273.529 compared to the mRNA-1273 vaccine.

Bivalent vaccines are one strategy to increase protection against currently circulating variants as well as broaden neutralization to previous and potentially yet-to-emerge variants^14,15^. When administered as a booster dose, the bivalent vaccine mRNA-1273.211 encoding for the Wuhan-1 and Beta (B.1.351) spike proteins induced neutralizing antibody responses in humans against B.1.351, Delta (B.1.617.2), and Omicron (BA.1) that were greater than that achieved by boosting with the parental mRNA-1273 vaccine^16,17^. Similarly, in interim data from other human studies, boosting with a bivalent mRNA-1273.214 vaccine targeting the Wuhan-1 and BA.1 strains elicited higher neutralizing antibody responses against BA.1, BA.2, and BA.4/5 than the mRNA-1273 booster, with neutralization of BA.4 and BA.5 assessed together, as the spike proteins of these two sub-lineages are the same^14,18^. Despite a lack of published data on the efficacy of bivalent Omicron-matched vaccines or boosters against infection by Omicron variants in humans, bivalent mRNA vaccine boosters that include Wuhan-1 and either BA.1 or BA.4/5 components recently were authorized in Europe and the United States, in part due to the urgent need to broaden protection against circulating SARS-CoV-2 variants.

Here, we evaluated in mice the antibody responses and protective activity against the prevailing circulating Omicron variant, BA.5, after a primary vaccination series or boosting with either of two Moderna bivalent vaccines, mRNA-1273.214 (containing 1:1 mix of mRNAs encoding Wuhan-1 and BA.1 spike proteins) and mRNA-1273.222 (1:1 mix of mRNAs encoding the Wuhan-1 and BA.4/5 spike proteins) and compared the results to monovalent vaccines that contain mRNAs encoding for a single spike antigen (Wuhan-1 [mRNA-1273], BA.1 [mRNA-1273.529], or BA.4/5 [mRNA1273.045]). In immunogenicity studies in BALB/c mice, performed in the context of a primary vaccination series, robust anti-spike antibody responses were detected with all mRNA vaccines, as measured against Wuhan-1 (S2P), BA.1 (S2P.529) and BA.4/5 (S2P.045) spike proteins. However, both bivalent vaccines induced broader neutralizing antibody responses than the constituent monovalent vaccines against pseudoviruses displaying Wuhan-1, BA.1, BA.2.75, or BA.4/5 spike proteins. In immunogenicity studies in K18-hACE2 mice performed seven months after a primary vaccination series with mRNA-1273, animals boosted with mRNA-1273.214 or mRNA-1273.222 had higher neutralization titers against authentic BA.1 and BA.5 viruses, as well as similar neutralization titers against Wuhan-1 and B.1.617.2 viruses, compared to animals boosted with mRNA-1273. This response correlated with increased protection one month later against challenge with BA.5, as the lowest viral RNA and pro-inflammatory cytokine levels in the lung were observed in mice administered mRNA-1273.214 or mRNA-1273.222 boosters. Thus, bivalent mRNA vaccine boosters that include mRNAs for both Wuhan-1 and Omicron spike proteins induce protective immunity against historical and current SARS-CoV-2 variant strains.

## RESULTS

### Preclinical bivalent Omicron-targeted mRNA vaccines induce robust antibody responses in BALB/c mice

Immunization with bivalent vaccines that include components targeting an Omicron spike and the original Wuhan-1 spikes could confer broader immunity. To begin to address this question, we generated two lipid-encapsulated (LNP) mRNA vaccines (mRNA-1273.529 and mRNA-1273.045) encoding a proline-stabilized SARS-CoV-2 spike from BA.1 and BA.4/5 viruses, respectively. The mRNA-LNPs then were combined with mRNA-1273 in a 1:1 ratio to generate bench-side mixed versions of mRNA-1273.214 and mRNA-1273.222. As a first test of their activity, we immunized BALB/c mice twice at a 3-week interval with 1 μg (total dose) of preclinical versions of mRNA-1273, mRNA-1273.529, mRNA-1273.045, mRNA-1273.214 or mRNA-1273.222 vaccines (**Fig 1A**). Three weeks after the first dose (day 21) and two weeks after the second dose (day 35), serum was analyzed for binding to Wuhan-1 (S2P), BA.1 (S2P.529), and BA.4/5 (S2P.045) spike proteins by ELISA (**Fig 1B**).

**Figure 1.**
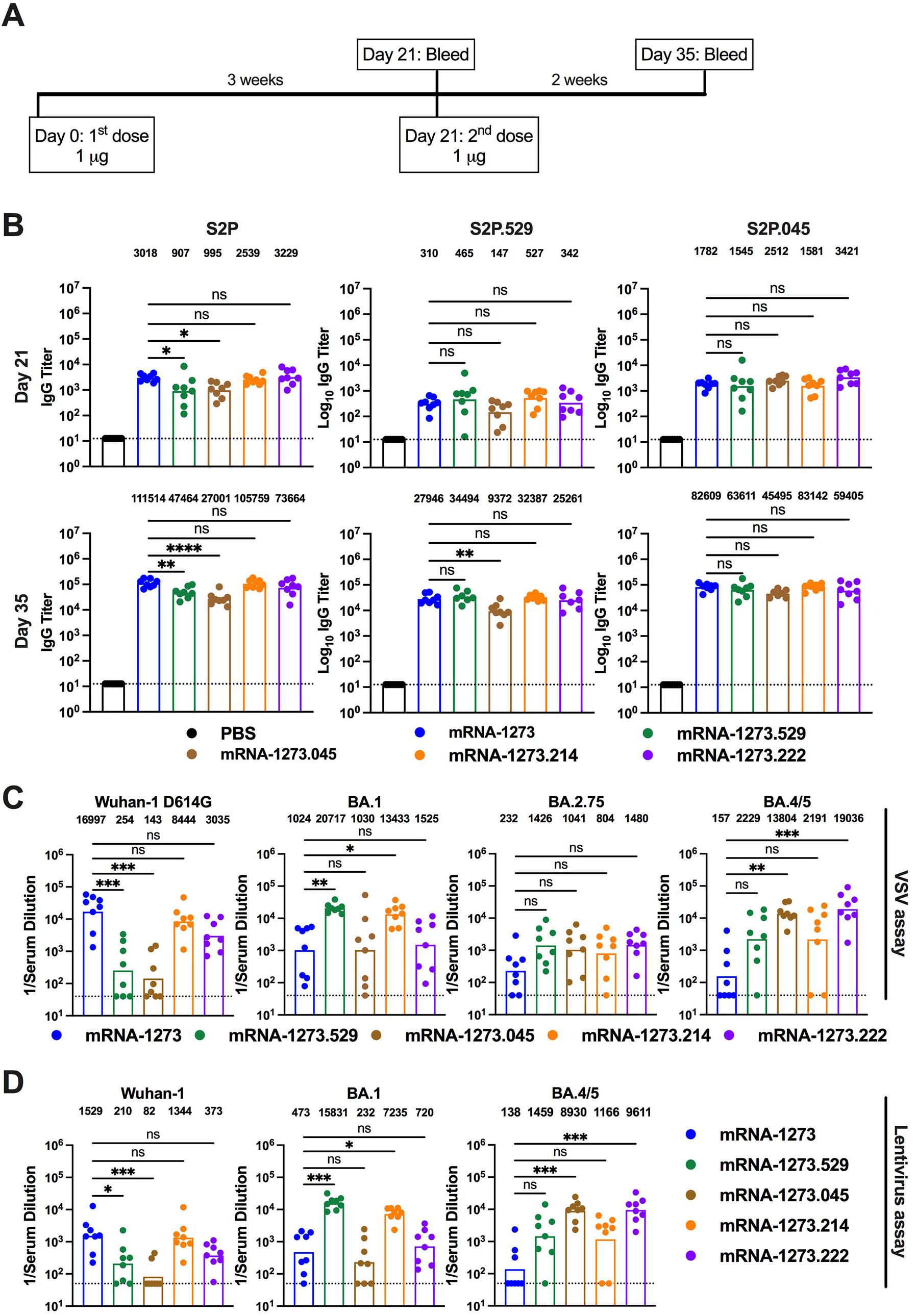
Robust antibody responses in BALB/c mice after a primary immunization series with preclinical versions of monovalent and bivalent mRNA vaccines. Six-to-eight-week-old female BALB/c mice were immunized twice over a three-week interval with PBS or 1 μg total dose of preclinical versions of mRNA-1273 [Wuhan-1 spike], mRNA-1273.529 [BA.1 spike], mRNA-1273.045 [BA.4/5 spike], mRNA-1273.214 [benchside 1:1 mixture of mRNA-1273 + mRNA-1273.529], or mRNA-1273.222 [benchside 1:1 mixture of mRNA-1273 + mRNA-1273.045]. Immediately before (day 21) or two weeks after (day 35) the second vaccine dose, serum was collected. **A**. Scheme of immunization and blood draws. **B**. Serum antibody binding to Wuhan-1 (S2P), BA.1 (S2P.529), or BA.4/5 (S2P.045) spike proteins by ELISA at Day 21 and Day 35 (n = 8, boxes illustrate mean values, dotted lines show the limit of detection [LOD]). **C**. Neutralizing activity of serum at day 35 against VSV pseudoviruses displaying the spike proteins of Wuhan-1 D614G, BA.1, BA.2.75, or BA.4/5 (n = 8, boxes illustrate geometric mean values, dotted lines show the LOD). GMT values are indicated above the columns. **D**. Neutralizing activity of serum at day 35 against pseudotyped lentiviruses displaying the spike proteins of Wuhan-1, BA.1, or BA.4/5 (n = 8, boxes illustrate geometric mean values, dotted lines show the LOD). GMT values are indicated above the columns. Statistical analysis. **B**. One-way ANOVA with Dunnett’s post-test. **C-D**. Kruskal-Wallis with Dunn’s post-test (ns, not significant; * *P* < 0.05; ** *P* < 0.01; *** *P* < 0.001; **** *P* < 0.0001).

Robust serum IgG binding was observed against S2P, S2P.529, and S2P.045 proteins after a two-dose primary series with monovalent mRNA-1273, mRNA-1273.529, and mRNA-1273.045 vaccines as well as bivalent mRNA-1273.214 and mRNA-1273.222 vaccines, compared to immunizing with PBS only. (a) <u>S2P</u>. On day 21, geometric mean titers (GMT) against S2P ranged from 907 to 3,229 and increased by approximately 23- to 52-fold on day 35, with values ranging from 27,001 to 111,514 across the vaccine groups (**Fig 1B**). On day 35, mice vaccinated with mRNA-1273, mRNA-1273.214, or mRNA-1273.222 achieved higher GMTs than mice vaccinated with mRNA-1273.529 or mRNA-1273.045 vaccines (b) <u>S2P.529</u>. Serum binding GMTs against S2P.529 at day 21 ranged from 147 to 527 and increased by approximately 61- to 90-fold on day 35, with values ranging from 9,372 to 34,494 across the vaccine groups (**Fig 1B**). There were no significant differences in binding titers against S2P.529 across most groups on day 35, although serum from mice vaccinated with mRNA-1273.045 showed reduced binding compared to mRNA-1273. (c) <u>S2P.045</u>. At day 21, serum IgG binding GMTs against S2P.045 ranged from 1,545 to 3,421 and increased by approximately 17- to 52-fold on day 35, with values ranging from 45,495 to 83,142 (**Fig 1B**). On day 35, robust IgG GMTs against S2P.045 were observed for all mRNA vaccinated groups with no substantive differences noted.

We next tested the inhibitory activity of serum antibodies from BALB/c mice that received two doses of the different preclinical mRNA vaccines using a vesicular stomatitis virus (VSV)-based neutralization assay with pseudoviruses displaying spike proteins of Wuhan-1 D614G, BA.1, BA.2.75, or BA.4/5 (**Fig 1C and Extended Data Fig 1**). Serum obtained at day 35 from mice vaccinated with mRNA-1273.222 or mRNA-1273.045 vaccines showed robust neutralizing antibody responses (GMT: 19,036 and 13,804, respectively) against BA.4/5. When compared to the neutralizing antibody response against Wuhan-1 D614G elicited by the mRNA-1273 vaccine (GMT: 16,997), the response against BA.4/5 elicited by mRNA-1273.222 vaccine was equivalent, if not slightly higher. Moreover, the neutralizing antibody response against Wuhan-1 D614G elicited by the mRNA-1273.222 vaccine was greater than that elicited by the mRNA-1273.045 vaccine (GMT: 3,035 and 143, respectively). As expected, serum from the mRNA-1273.214 and mRNA-1273.529 vaccine recipients robustly inhibited infection of BA.1 pseudoviruses, with slightly greater titers elicited by the mRNA-1273.529 vaccine (GMT: 13,433 and 20,717, respectively). However, the mRNA-1273.214 vaccine induced much greater serum neutralizing activity against Wuhan-1 D614G (GMT: 8,443) than the mRNA-1273.529 vaccine (GMT:196). The mRNA-1273 vaccine showed a robust response against Wuhan-1 D614G (GMT: 16,997), but less effective responses against BA.1 (GMT: 1,024), BA.2.75 (GMT: 232), or BA.4/5 (GMT:157). All of the Omicron-matched vaccines induced slightly greater (3.5 to 6.4-fold) neutralizing antibody responses than mRNA-1273 against BA.2.75, although these differences did not attain statistical significance.

We also evaluated serum neutralizing antibody capacity using a lentivirus-based pseudovirus assay with virions displaying Wuhan-1, BA.1, or BA.4/5 spike proteins. The neutralizing antibody responses measured with lentiviruses were similar to those obtained using VSV pseudoviruses (**Fig 1D and Extended Data Fig 2**). On day 35, mice immunized with the mRNA-1273.045 or mRNA-1273.222 vaccines had robust responses against BA.4/5 (GMT: 1,166 and 9,611, respectively), although the responses against Wuhan-1 and BA.1 were lower. Serum from animals immunized with mRNA-1273 vaccine efficiently neutralized Wuhan-1 (GMT: 1,529) but the responses against BA.1 and BA.4/5 were lower (GMT: 473 and 138, respectively). The mRNA-1273.529 and mRNA-1273.214 vaccines induced strong neutralizing antibody responses against BA.1 (GMT: 15,831 and 7,235, respectively) with less inhibitory activity against Wuhan-1 (GMT: 210 and 1,344, respectively) and BA.4/5 (GMT: 1,459 and 1,166, respectively). Overall, based on data from the VSV and lentivirus pseudovirus assays, both bivalent mRNA-1273.222 and mRNA-1273.214 vaccines offered the most neutralization breadth.

### Clinically representative versions of bivalent mRNA vaccines induce robust neutralizing antibody responses in BALB/c mice

We next evaluated the immunogenicity of clinically representative versions of mRNA-1273.214 and mRNA-1273.222, where two monovalent mRNAs were separately formulated into LNPs and then mixed in a 1:1 ratio in the vial, a process that is representative of the commercial drug product. These versions were compared to responses obtained with mRNA-1273. BALB/c mice were immunized twice at a 3-week interval with 1 μg (total dose) of mRNA-1273, mRNA-1273.214, or mRNA-1273.222 vaccines (**Fig 2A**). Two weeks after the second dose (day 35), serum was collected and analyzed for neutralizing activity using VSV-based pseudoviruses displaying Wuhan-1 D614G, BA.1, BA.2.75, or BA.4/5 spike proteins (**Fig 2B and Extended Data Fig 3**). Whereas mRNA-1273 induced a robust neutralizing antibody response against Wuhan-1 D614G (GMT: 28,920), 30 to 194-fold less activity was measured against pseudoviruses displaying BA.1, BA.2.75, and BA.4/5. The breadth of neutralizing activity seen with the bivalent mRNA-1273.214 vaccine was better, with the greatest responses against the matched BA.1 (GMT: 13,183) and 1.7 to 52-fold reductions against Wuhan-1 D614G, BA.2.75, and BA.4/5, with the lowest potency against BA.4/5 (GMT: 293). The mRNA-1273.222 vaccine achieved the broadest inhibitory activity with the highest neutralizing titers against the matched BA.4/5 (GMT: 15,561) and only 1.7 to 6.7-fold reductions in activity against Wuhan-1 D614G, BA.1, and BA.2.75.

**Figure 2.**
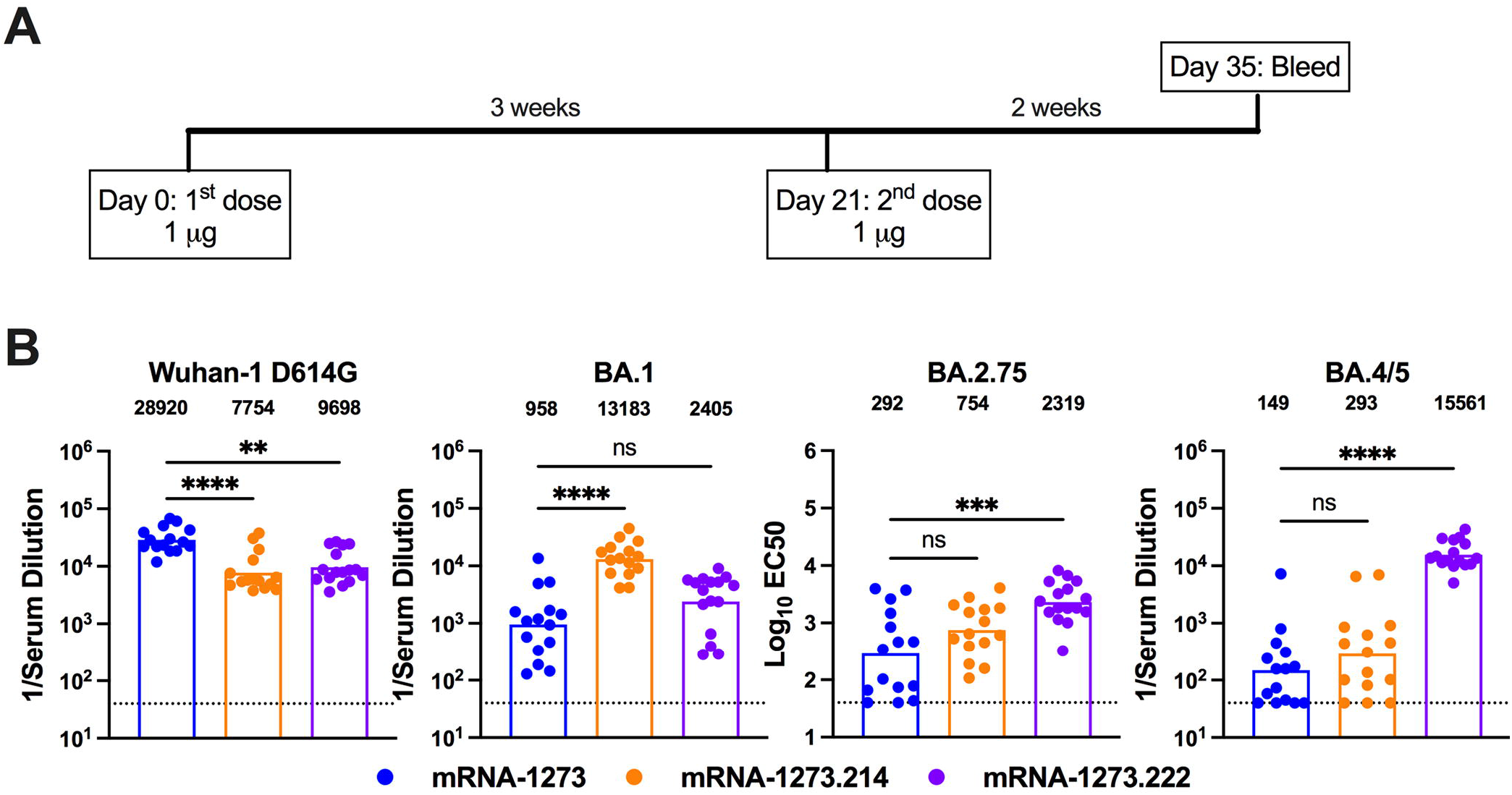
Robust neutralizing antibody responses in BALB/c mice after primary series immunization with clinically representative versions of mRNA-1273, mRNA-1273.214, and mRNA-1273.222. Six-to-eight-week-old female BALB/c mice were immunized twice over a three-week interval with PBS or 1 μg total dose of clinically representative versions of mRNA-1273, mRNA-1273.214 [1::1 mixture in the vial of separately formulated mRNA-1273 and mRNA-1273.529], or mRNA-1273.222 [1:1 mixture in the vial of separately formulated mRNA-1273 and mRNA-1273.045]. Immediately before (day 21) or two weeks after (day 35) the second vaccine dose, serum was collected. **A**. Scheme of immunization and blood draws. **B**. Neutralizing activity of serum at day 35 against VSV pseudoviruses displaying the spike proteins of Wuhan-1 D614G, BA.1, BA.2.75, or BA.4/5 (n = 16, boxes illustrate geometric mean values, dotted lines show the LOD). GMT values are indicated above the columns. Statistical analysis. Kruskal-Wallis with Dunn’s post-test (ns, not significant; ** *P* < 0.01; **** *P* < 0.0001).

### Boosting with clinically representative versions of bivalent mRNA vaccines enhances neutralizing antibody responses against Omicron variants and confers protection against BA.**5 infection in mice**

We next evaluated the performance of the bivalent vaccines as booster injections in mice, as mRNA-1273.214 and mRNA-1273.222 have been authorized as boosters in SARS-CoV-2 antigen-experienced humans. We took advantage of an existing cohort of female K18-hACE2 transgenic C57BL/6 mice that had received two 0.25 μg doses of mRNA-1273 or control mRNA vaccine over a three-week interval and were rested subsequently for 31 weeks (**Fig 3A**). The 0.25 μg dose of mRNA vaccine was used because the B and T cell responses generated in C57BL/6 mice with this dose approximate in magnitude those observed in humans receiving 100 μg doses^12,19^. Blood was collected (pre-boost sample), and groups of animals were boosted with either PBS (sham control), or 0.25 μg of control mRNA, mRNA-1273, mRNA-1273.214, or mRNA-1273.222 vaccines. Four weeks later, a post-boost blood sample was collected (**Fig 3A**), and the neutralizing activity of pre- and post-boost serum antibodies was determined using authentic SARS-CoV-2 viruses. At 31 weeks after completion of the primary mRNA-1273 vaccination series, pre-boost neutralizing antibody levels against WA1/2020 D614G (GMT: 454) and B.1.617.2 (GMT: 277) were above the expected threshold (∼1/60) of protection^20^ (**Fig 3B, Extended Data Figs 4 and 5**). However, these samples showed less or no neutralizing activity (GMT: 63) against BA.1 or BA.5 at the lowest dilution tested (**Fig 3B, Extended Data Figs 4 and 5**), consistent with a 20-fold reduction reported in serum samples from humans immunized with mRNA vaccines targeting ancestral SARS-CoV-2 strains^6,7^.

**Figure 3.**
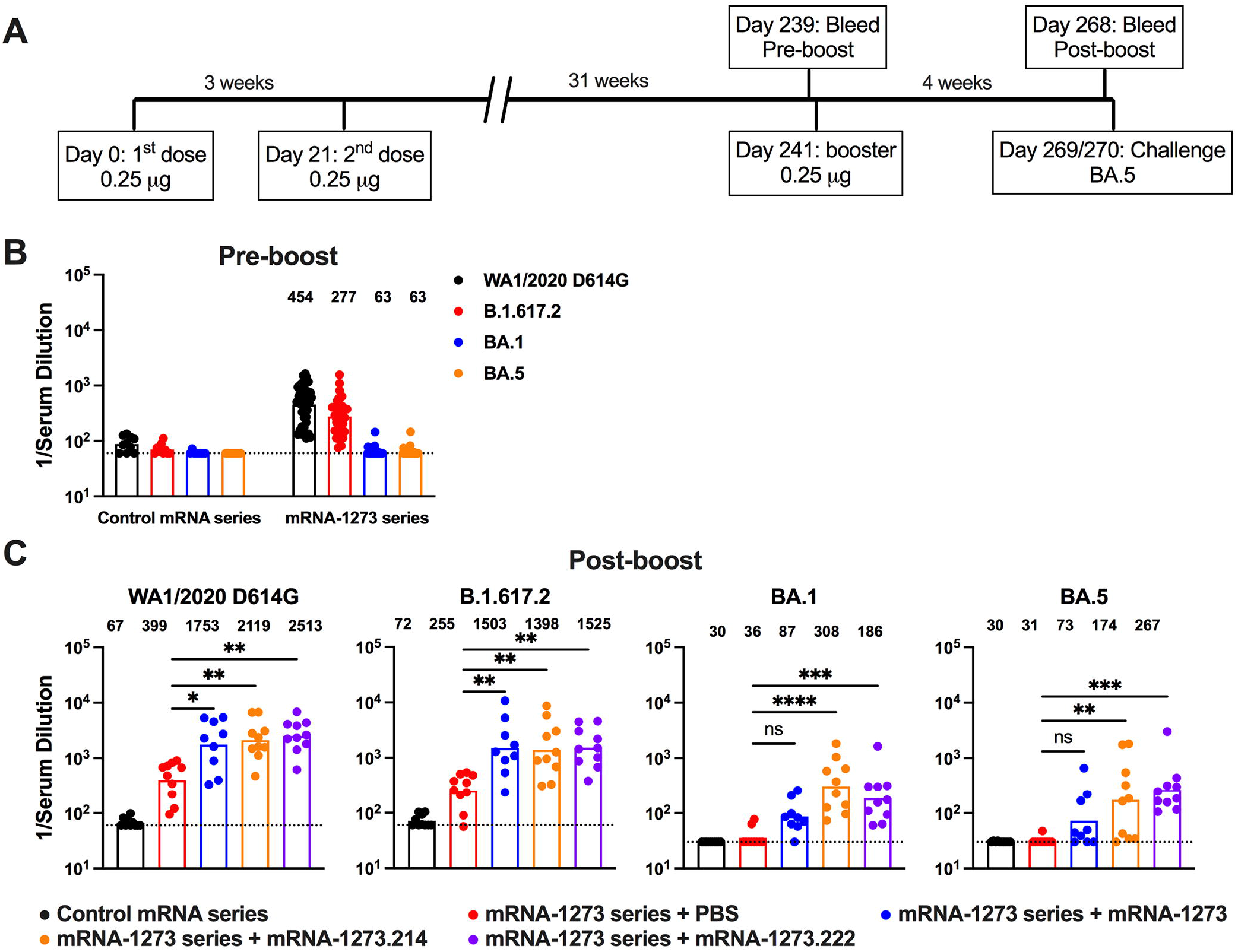
Neutralizing antibody responses in K18-hACE2 mice after boosting with clinically representative versions of mRNA-1273, mRNA-1273.214, and mRNA-1273.222. Seven-week-old female K18-hACE2 mice were immunized with 0.25 μg of control mRNA or mRNA-1273 vaccine and then boosted 31 weeks later with PBS, 0.25 μg of control mRNA, or 0.25 μg of clinically representative versions of mRNA-1273, mRNA-1273.214, or mRNA-1273.222 vaccines. **A**. Scheme of immunizations, blood draws and virus challenge. **B-C**. Serum neutralizing antibody responses immediately before (**B**, pre-boost) and four weeks after (**C**, post-boost) receiving the indicated mRNA boosters or PBS as judged by focus reduction neutralization test (FRNT) with authentic WA1/2020 D614G, B.1.617.2, BA.1, and BA.5 viruses (n = 9-10, two experiments, boxes illustrate geometric mean values, GMT values are indicated at the top of the graphs, dotted lines show the LOD). Statistical analysis. **C**: Kruskal-Wallis with Dunn’s post-test, ns, not significant; * *P* < 0.05; ** *P* < 0.01; *** *P* < 0.001; **** *P* < 0.0001).

Four weeks after boosting with mRNA-1273, mRNA-1273.214, or mRNA-1273.222, neutralizing titers against WA1/2020 D614G and B.1.617.2 were approximately 4.2 to 6.0-fold and 4.8 to 5.5-fold higher, respectively, than before boosting (**Fig 3C, Extended Data Figs 4-6**). Boosting with mRNA-1273.214 or mRNA-1273.222 resulted in increased neutralizing titers against BA.1 (3 to 5.1-fold) and BA.5 (2.6 to 4.4-fold), respectively (**Fig 3C, Extended Data Figs 4-6**), whereas mRNA-1273 boosted titers by a lesser degree (1.0 to 1.2-fold). Thus, both Omicron-matched bivalent boosters augmented serum neutralizing activity against BA.1 and BA.5 more than the parental mRNA-1273 booster.

One or two days after the post-boost bleed, K18-hACE2 transgenic mice were challenged by the intranasal route with 10^4^ FFU of BA.5 (**Fig 3A**), and at 4 days post-infection (dpi) viral RNA levels were measured in the nasal washes, nasal turbinates, and lungs (**Fig 4A**). Although Omicron strains are less pathogenic in mice and do not cause weight loss or mortality^21,22^, viral replication occurs allowing for evaluation of vaccine protection. In the upper respiratory tract (nasal turbinates and nasal washes), mice boosted with PBS or with mRNA-1273, mRNA-1273-214, or mRNA-1273.222 vaccines showed similarly reduced levels of BA.5 viral RNA at 4 dpi in comparison to animals administered the control vaccine (**Fig 4A**). However, mice immunized with two doses of mRNA-1273 and boosted with PBS sustained levels of viral RNA in the lungs that were only slightly less than the control vaccine, suggesting that a primary mRNA-1273 vaccination series provides limited protection against lower respiratory tract infection by BA.5. In contrast, mRNA-1273, mRNA-1273-214, or mRNA-1273.222 vaccines showed greater protection against BA.5 infection in the lung with 1,374 to 28,436-fold reductions in viral RNA (**Fig 4A**). Moreover, boosting with the bivalent vaccines resulted in lower (7 to 21-fold) viral RNA levels in the lungs than the mRNA-1273 vaccine. Analysis of infectious virus in the lung at 4 dpi using plaque assays showed substantial reductions in viral burden in animals boosted with mRNA-1273, mRNA-1273-214, or mRNA-1273.222 vaccines compared to those receiving a control vaccine or immunized with two doses of mRNA-1273 and boosted with PBS (**Fig 4B**).

**Figure 4.**
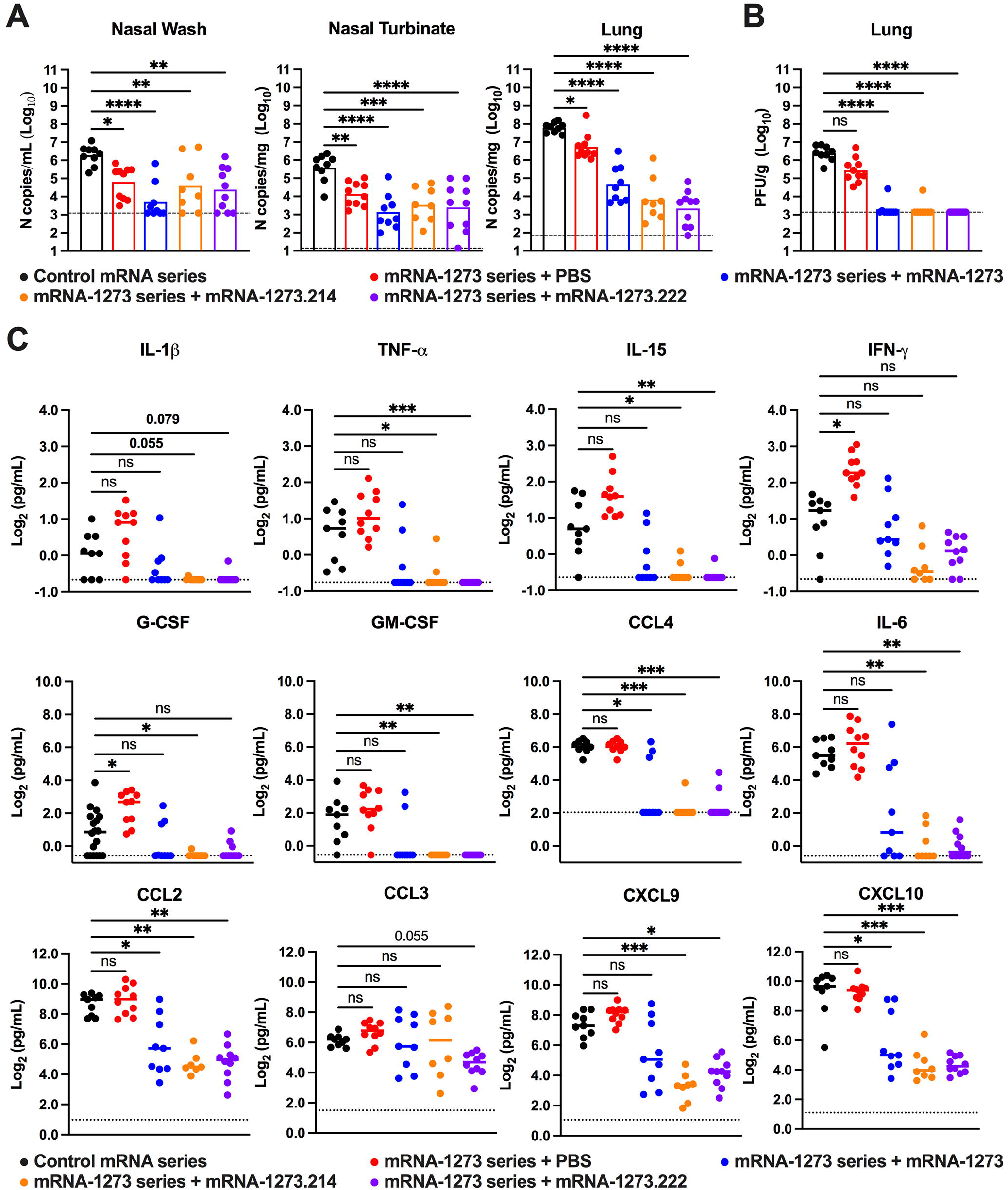
Protection of K18-hACE2 mice from BA.5 challenge after boosting with clinically representative versions of mRNA-1273, mRNA-1273.214, and mRNA-1273.222. Seven-week-old female K18-hACE2 mice were immunized with 0.25 μg of control mRNA or mRNA-1273, boosted 31 weeks later with PBS, 0.25 μg of control mRNA, or 0.25 μg of clinically representative versions of mRNA-1273, mRNA-1273.214, or mRNA-1273.222 vaccines, and then one month later challenged via intranasal route with 10^4^ focus-forming units (FFU) of BA.5. **A**. Viral RNA levels at 4 dpi in the nasal washes, nasal turbinates, and lungs (n = 8-10 per group, two experiments, boxes illustrate mean values, dotted lines show LOD; One-way ANOVA with Dunnett’s post-test: * *P* < 0.05; ** *P* < 0.01; *** *P* < 0.001; **** *P* < 0.0001). **B**. Infectious viral load at 4 dpi in the lungs after BA.5 challenge of vaccinated and boosted mice as determined by plaque assay (n= 8-10 per group, two experiments, boxes illustrate mean values, dotted lines show LOD; Kruskal Wallis with Dunn’s post-test: ns, *P* > 0.05; **** *P* < 0.0001). **C**. Cytokine and chemokine levels in lung homogenates at 4 dpi. Data are expressed as fold-change relative to naive mice, and log_2_ values are plotted (n = 8-10 per group except naïve where n = 4, two experiments, lines illustrates median values, dotted lines indicate LOD for each respective analyte based on standard curves; one-way Kruskal Wallis ANOVA with Dunn’s post-test: ns, *P* > 0.05; * *P* < 0.05; ** *P* < 0.01; *** *P* < 0.001; **** *P* < 0.0001). The absolute values are shown in **Supplementary Table S1**.

As another gauge of vaccine-induced protection, we measured cytokine and chemokine levels in the lung of the BA.5-challenged K18-hACE2 mice at 4 dpi using a multiplexed assay (**Fig 4C and Supplementary Table S1**). Mice immunized with the control vaccine or those receiving a primary vaccination series with mRNA-1273 and a booster dose of PBS showed higher levels of many inflammatory cytokines and chemokines in lung homogenates than unvaccinated, unchallenged (naïve) animals. In comparison, substantially lower or undetectable levels of pro-inflammatory cytokines and chemokines were detected in the lungs of BA.5-challenged mice that were boosted with mRNA-1273, mRNA-1273-214, or mRNA-1273.222 vaccines. While there was some variability in the data, for several cytokines and chemokines (*e.g*., IFN-γ, CCL2, CCL3, and CXCL9), levels trended lower after boosting with mRNA-1273-214 and mRNA-1273.222 compared to mRNA-1273. Thus, and consistent with the virological data, protection against BA.5-induced lung inflammation was significantly increased by boosting with mRNA vaccines, with a modest improvement observed after boosting with bivalent compared to the monovalent mRNA vaccines.

## DISCUSSION

Vaccine-induced immunity against SARS-CoV-2 has reduced human disease and curtailed the COVID-19 pandemic. However, the emergence of SARS-CoV-2 variants with constellations of amino acid changes in regions of the spike protein that bind neutralizing antibodies jeopardizes immunity derived from vaccines designed against the historical Wuhan-1 SARS-CoV-2 strain. The objective of this study was to evaluate in mice the activity of pre-clinical and clinically representative Omicron-matched (BA.1 or BA.4/5) bivalent mRNA vaccines when administered as a primary vaccination series or booster dose and to assess the breadth of neutralizing antibody responses and ability to protect against currently circulating Omicron variants. The animal studies performed here support the use of bivalent mRNA-1273.214 and mRNA-1273.222 vaccines that were recently authorized for Europe and the United States, respectively.

We first compared the immunogenicity of the bivalent and constituent monovalent mRNA vaccines in BALB/c mice in the context of a two-dose primary immunization series. Although bivalent mRNA vaccines are conceived principally as boosters since most of the global population has been previously infected or vaccinated, the analysis of the antibody response after a primary immunization series provides insight into the potential breadth. Robust serum antibody binding responses were detected against S2P, S2P.529, and S2P.045 proteins by all vaccines, although the mRNA-1273.529 and mRNA-1273.045 vaccines had lower titers against non-matched spike antigens. Moreover, serum generated from the bivalent mRNA-1273.222 and mRNA-1273.214 vaccines potently neutralized infection of both Omicron (BA.1. and BA.4/5) pseudoviruses, as well as those displaying the historical Wuhan-1 D614G spike, demonstrating the best neutralization breadth. These results are consistent with studies showing that monovalent vaccines that match the spike protein generate greater inhibitory responses against specific variants relative to historical viruses^12,23,24^. In the context of the booster studies, the bivalent BA.1 and BA.4/5-matched vaccines induced serum antibody responses that more broadly neutralized infection of several authentic viruses, including WA1/2020 D614G, B.1.617.2, BA.1, and BA.5. These results are consistent with human serum data obtained after immunization with bivalent mRNA vaccines targeting B.1.351^16,17^ or BA.1^14,18^. Increases in neutralizing antibody breadth with bivalent vaccine formulations or boosters also have been reported in the context of inactivated^25,26^ and spike protein-based^27-29^ SARS-CoV-2 vaccine candidates.

Seven months after completion of a primary mRNA-1273 vaccination series, K18-hACE2 mice were boosted with PBS, mRNA-1273, mRNA-1273.214, or mRNA-1273.222, and then one month later challenged with BA.5. We used lower priming and boosting vaccine doses (0.25 μg per mouse) than in our BALB/c mice studies, since we had observed that the levels of immunity induced by the 0.25 μg dose in K18-hACE2 mice more closely match what is seen in humans^12,19^. For example, seven months after receiving two 0.25 μg doses of mRNA-1273, moderately high (1/454 and 1/277) neutralizing titers still were present against the matched WA1/2020 D614G and more closely related B.1.617.2, respectively, in K18-hACE2 mice, but this inhibitory activity was almost completely lost against BA.1 and BA.5, similar to that seen in humans^6,7^. Compared to a third dose of mRNA-1273, both bivalent mRNA-1273.214 and mRNA-1273.222 vaccine boosters induced greater neutralizing antibody responses against BA.1 and BA.5, and this correlated with slightly increased protection against infection and inflammation in the lung after intranasal challenge with BA.5 compared to the parental mRNA-1273 boost. In comparison, animals that received a primary mRNA-1273 series, and were boosted with PBS, showed rather marginal protection against lung infection. These results showing protective benefit of matched bivalent vaccine boosters targeting Omicron variants are consistent with studies in mice immunized with mRNA-1273, boosted with a monovalent mRNA-1273.529 vaccine, and challenged with BA.1^12^, and predictive models in humans^30^.

Notwithstanding the increased immunogenicity and protection by the bivalent vaccines, boosting with the parental mRNA-1273 vaccine alone conferred protection against infection (119-fold reduction in viral RNA levels and 142-fold reduction in infectious virus compared to boosting with PBS) and inflammation in the lung against BA.5 despite inducing rather limited levels of serum neutralizing antibodies against this variant. These findings are consistent with studies in non-human primates^13^ and could reflect effects of neutralizing antibodies below the limit of our assay detection (<1/30), non-neutralizing, cross-reactive antibodies against BA.5 that promote clearance through Fc effector function activities^31,32^, cross-reactive T cell responses^33,34^, or anamnestic B cell responses that rapidly generate cross-reactive neutralizing antibodies. Apart from this, our experiments show that two bivalent mRNA vaccines including components against BA.1 or BA.4/5 had relatively equivalent protective effects against BA.5 in the lungs. Although there is a trend towards lower levels of BA.5 RNA after boosting with mRNA-1273.222 compared to mRNA-1273.214 vaccines, our studies were not powered sufficiently to establish this increased protection, and larger cohorts would be needed to reach this conclusion.

All mRNA vaccine boosters conferred protection in the upper respiratory tract, with reductions in viral RNA levels measured in the nasal washes and nasal turbinates at 4 dpi. The bivalent and mRNA-1273 vaccine boosters performed equivalently, with similar reductions in viral burden compared to the control vaccine. It is unclear why the differences in protection in the lung between bivalent and parental monovalent mRNA vaccine boosters did not extend to the nasal washes and turbinates, although it may be because neutralizing IgG poorly penetrates this compartment ^35^, and immune protection in the upper respiratory is mediated by other components (*e.g*., T cells or trained innate immunity) not assayed here. Regardless, our data showing that both bivalent vaccine boosters confer increased neutralizing activity as well as protection in the lungs against BA.5 supports the recent decision for roll-out of BA.1 or BA.4/5-based bivalent boosters.

### Limitations of study

We note several limitations in our study. (1) Female BALB/c and K18-hACE2 mice were used to allow for group caging. Follow-up experiments with male mice and larger cohorts are needed to extend these results and possibly detect differences between mRNA-1273.214 and mRNA-1273.222 boosters. (2) We challenged K18-hACE2 mice with a BA.5 isolate. While BA.5 is currently the dominant circulating strain (reaching up to ∼89% in the United States (https://covid.cdc.gov/covid-data-tracker/#variant-proportions) for the week ending September 3, 2022), infection experiments using BA.2.75, BA.4.6, or other emerging strains may be informative to determine the breadth of protection. (3) Our analysis did not account for non-neutralizing antibody or cross-reactive T cell responses, both of which could impact protective immunity. (4) We analyzed protection in the lung one month after boosting. A time course analysis is needed to assess the durability of the boosted immune response. (5) Experiments were performed in mice to allow for rapid testing and multiple comparison groups. Vaccination, boosting, and BA.5 challenge in other animal models (*e.g*., hamsters and non-human primate) and ultimately in humans is required for corroboration. (6) While our studies evaluated differences in breadth of serum neutralizing antibody responses, a repertoire analysis at the monoclonal level could provide insight as to how bivalent mRNA vaccines inhibit variant strains.

## Acknowledgements

This study was supported by the NIH (R01 AI157155, NIAID Centers of Excellence for Influenza Research and Response (CEIRR) contracts HHSN272201400008C, 75N93021C00014, and 75N93019C00051). We thank Mehul Suthar for the BA.5 isolate used in this study. We also acknowledge and thank Michael Whitt for support on VSV-based pseudovirus production.

## Author contributions

G.-Y.C., G.S.-J., and A.N. performed variant monitoring and Omicron-variant vaccine design and quality control. S.M.S. performed and analyzed authentic virus neutralization assays. S.M.S., B.W. and B.Y. performed mouse experiments. B.W. and B.Y. performed and analyzed viral burden analyses. B.Y. analyzed chemokine and cytokine data. H.J., P.M., and N.J.A. performed ELISA binding experiments and analysis. K.W., D.L., D.M.B., and L.A. performed VSV-pseudovirus neutralization assays and analysis. S.D.S., S.O., R.A.K., and N.D.-R. performed lentivirus pseudovirus neutralization assays and analysis. A.C., S.E., and D.K.E. provided mRNA vaccines and helped to design experiments. L.B.T. and M.S.D. designed studies and supervised the research. M.S.D., L.B.T., and D.K.E. wrote the initial draft, with the other authors providing editorial comments.

## EXTENDED DATA FIGURE LEGENDS

**Extended Data Figure 1. Comparison of serum neutralization using VSV pseudoviruses expressing Wuhan-1 D614G, BA.1, BA.2.75, or BA.4/5 spike proteins, Related to Figure 1**. BALB/c mice were immunized with two 1 μg doses of preclinical versions of mRNA-1273, mRNA-1273.529, mRNA-1273.045, mRNA-1273.214 or mRNA-1273.222 vaccines. Serum neutralizing antibody responses against Wuhan-1 D614G, BA.1, and BA.4/5 were assessed two weeks after the second dose using VSV pseudoviruses. Representative neutralization curves (n = 2) corresponding to individual mice are shown for the indicated vaccines.

**Extended Data Figure 2. Comparison of serum neutralization using pseudotyped lentiviruses expressing Wuhan-1, BA.1, or BA.4/5 spike proteins, Related to Figure 1**. BALB/c mice were immunized with two 1 μg doses of preclinical versions of mRNA-1273, mRNA-1273.529, mRNA-1273.045, mRNA-1273.214 or mRNA-1273.222 vaccines. Serum neutralizing antibody responses against Wuhan-1, BA.1, and BA.4/5 were assessed two weeks after the second dose using pseudotyped lentiviruses. Average neutralization curves (n = 2) corresponding to individual mice are shown for the indicated vaccines.

**Extended Data Figure 3. Comparison of serum neutralization using VSV pseudoviruses expressing Wuhan-1 D614G, BA.1, BA.2.75, or BA.4/5 spike proteins, Related to Fig 2ure**. BALB/c mice were immunized with two 1 μg doses of clinically representative versions of mRNA-1273, mRNA-1273.214 or mRNA-1273.222 vaccines. Serum neutralizing antibody responses against Wuhan-1 D614G, BA.1, BA.2.75, and BA.4/5 were assessed two weeks after the second dose using VSV pseudoviruses. Representative neutralization curves (n = 2) corresponding to individual mice are shown for the indicated vaccines.

**Extended Data Figure 4. Comparison of serum neutralization of authentic WA1/2020 D614G, B.1.617.2, BA.1, and BA.5 viruses before and after boosting, Related to Figure 3**. Seven-week-old female K18-hACE2 mice were immunized with two sequential 0.25 μg doses of control mRNA or mRNA-1273 and then boosted 31 weeks later with PBS, 0.25 μg of control mRNA, or 0.25 μg mRNA-1273, mRNA-1273.214, or mRNA-1273.222. Paired analysis of pre- and post-boost serum neutralizing titers against WA1/2020 D614G, B.1.617.2, BA.1 and BA.5 from samples obtained from animals (data from **Figure 3**) (n = 9-10, two experiments). GMT values are indicated at the top of the graphs. Statistical analysis: Wilcoxon signed-rank test (ns, not significant; ** *P* < 0.01).

**Extended Data Figure 5. Pre-boost serum neutralization of authentic WA1/2020 D614G, B.1.617.2, BA.1, and BA.5 viruses, Related to Figure 3**. Seven-week-old female K18-hACE2 mice were immunized with two sequential 0.25 μg doses of control mRNA or mRNA-1273 and then boosted 31 weeks later with PBS, 0.25 μg of control mRNA, or 0.25 μg mRNA-1273, mRNA-1273.214, or mRNA-1273.222. Neutralizing antibody responses against WA1/2020 D614G, B.1.617.2, BA.1, and BA.5 from serum immediately before boosting with the indicated vaccines. Neutralization curves (FRNT analysis) corresponding to individual mice are shown for the indicated immunizations. Serum are from two independent experiments, and each point represents the mean of two technical replicates.

**Extended Data Figure 6. Post-boost serum neutralization of authentic WA1/2020 D614G, B.1.617.2, BA.1, and BA.5 viruses, Related to Figure 3**. Seven-week-old female K18-hACE2 mice were immunized with two sequential 0.25 μg doses of control mRNA or mRNA-1273 and then boosted 31 weeks later with PBS, 0.25 μg of control mRNA, or 0.25 μg mRNA-1273, mRNA-1273.214, or mRNA-1273.222. Neutralizing antibody responses against WA1/2020 D614G, B.1.617.2, BA.1, and BA.5 from serum one month after boosting with the indicated vaccines. Neutralization curves (FRNT analysis) corresponding to individual mice are shown for the indicated immunizations. Serum are from two independent experiments, and each point represents the mean of two technical replicates.

## SUPPLEMENTAL TABLE

**Supplemental Table S1. Cytokine and chemokine concentrations in the lungs of immunized K18-hACE2 mice challenged with BA.5, Related to Figure 4**.

## METHODS

### Cells

African green monkey Vero-TMPRSS2^36^ and Vero-hACE2-TMPRRS2^37^ cells were cultured at 37°C in Dulbecco’s Modified Eagle medium (DMEM) supplemented with 10% fetal bovine serum (FBS), 10 mM HEPES pH 7.3, 1 mM sodium pyruvate, 1× non-essential amino acids, and 100 U/mL of penicillin–streptomycin. Vero-TMPRSS2 cells were supplemented with 5 µg/mL of blasticidin. Vero-hACE2-TMPRSS2 cells were supplemented with 10 µg/mL of puromycin. All cells routinely tested negative for mycoplasma using a PCR-based assay.

### Viruses

The WA1/2020 D614G and B.1.617.2 strains were described previously^19,38^. The BA.1 isolate (hCoV-19/USA/WI-WSLH-221686/2021) was obtained from an individual in Wisconsin as a mid-turbinate nasal swab^39^. The BA.5 isolate was isolated in California (hCoV-19/USA/CA-Stanford-79_S31/2022) and a gift of M. Suthar (Emory University). All viruses were passaged once on Vero-TMPRSS2 cells and subjected to next-generation sequencing^37^ to confirm the introduction and stability of substitutions. All virus experiments were performed in an approved biosafety level 3 (BSL-3) facility.

### Mice

Animal studies were carried out in accordance with the recommendations in the Guide for the Care and Use of Laboratory Animals of the National Institutes of Health. For studies (K18-hACE2 mice) at Washington University School of Medicine, the protocols were approved by the Institutional Animal Care and Use Committee at the Washington University School of Medicine (assurance number A3381–01). Virus inoculations were performed under anesthesia that was induced and maintained with ketamine hydrochloride and xylazine, and all efforts were made to minimize animal suffering. For studies with BALB/c mice, animal experiments were carried out in compliance with approval from the Animal Care and Use Committee of Moderna, Inc. Sample size for animal experiments was determined on the basis of criteria set by the institutional Animal Care and Use Committee. Experiments were neither randomized nor blinded.

Heterozygous K18-hACE2 C57BL/6J mice (strain: 2B6.Cg-Tg(K18-ACE2)2Prlmn/J, Cat # 34860) were obtained from The Jackson Laboratory. BALB/c mice (strain: BALB/cAnNCrl, Cat # 028) were obtained from Charles River Laboratories. Animals were housed in groups and fed standard chow diets.

### mRNA vaccine and lipid nanoparticle production process

A sequence-optimized mRNA encoding prefusion-stabilized Wuhan-Hu-1 (mRNA-1273), BA.1 (mRNA-1273.529, the Omicron gene component in mRNA-1273.214), and BA.5 (mRNA-1273.045, the Omicron gene component in mRNA-1273.222) The genes of SARS-CoV-2 S2P proteins were synthesized *in vitro* using an optimized T7 RNA polymerase-mediated transcription reaction with complete replacement of uridine by N1m-pseudouridine^40^. In addition to the two proline substitution, the BA.1 spike gene in the mRNA-1273.529 and mRNA-1273.214 vaccines encoded the following substitutions: A67V, Δ69-70, T95I, G142D, Δ143-145, Δ211, L212I, ins214EPE, G339D, S371L, S373P, S375F, K417N, N440K, G446S, S477N, T478K, E484A, Q493R, G496S, Q498R, N501Y, Y505H, T547K, D614G, H655Y, N679K, P681H, N764K, D796Y, N856K, Q954H, N969K, and L981F. The BA.5 gene in mRNA-1273.045 and mRNA-222 vaccines encoded the following substitutions: T19I, Δ24-26, A27S, Δ69-70, G142D, V213G, G339D, S371F, S373P, S375F, T376A, D405N, R408S, K417N, N440K, L452R, S477N, T478K, E484A, F486V, Q498R, N501Y, Y505H, D614G, H655Y, N679K, P681H, N764K, D796Y, Q954H, N969K.

A non-translating control mRNA was synthesized and formulated into lipid nanoparticles as previously described^41^. The reaction included a DNA template containing the immunogen open-reading frame flanked by 5’ untranslated region (UTR) and 3’ UTR sequences and was terminated by an encoded polyA tail. After RNA transcription, the cap-1 structure was added using the vaccinia virus capping enzyme and 2’-*O*-methyltransferase (New England Biolabs). The mRNA was purified by oligo-dT affinity purification, buffer exchanged by tangential flow filtration into sodium acetate, pH 5.0, sterile filtered, and kept frozen at –20°C until further use.

The mRNA was encapsulated in a lipid nanoparticle through a modified ethanol-drop nanoprecipitation process described previously^42^. Ionizable, structural, helper, and polyethylene glycol lipids were briefly mixed with mRNA in an acetate buffer, pH 5.0, at a ratio of 2.5:1 (lipid:mRNA). The mixture was neutralized with Tris-HCl, pH 7.5, sucrose was added as a cryoprotectant, and the final solution was sterile-filtered. Vials were filled with formulated lipid nanoparticle and stored frozen at –20°C until further use. The vaccine product underwent analytical characterization, which included the determination of particle size and polydispersity, encapsulation, mRNA purity, double-stranded RNA content, osmolality, pH, endotoxin, and bioburden, and the material was deemed acceptable for *in vivo* study. The preclinical material used in this study were: (1) monovalent mRNA-1273 vaccine that contains a single mRNA encoding the SARS-CoV-2 S2P antigen; (2) monovalent mRNA-1273.529 vaccine that contains a single mRNA encoding the SARS-CoV-2 S2P antigen for BA.1; (3) monovalent mRNA-1273.045 vaccine that contains a single mRNA encoding the SARS-CoV-2 S2P antigen of the BA.4/BA.5 subvariants of Omicron; (4) research-grade bivalent mRNA-1273.214 vaccine, which is a 1:1 bench side mix of separately formulated mRNA-1273 and mRNA-1273.529 vaccines; and (5) research grade bivalent mRNA-1273.222 vaccine, which is a 1:1 bench side mix of separately formulated mRNA-1273 and mRNA-1273.045 vaccines; (6) clinically representative bivalent mRNA-1273.214 vaccine, which is a 1:1 mix in the vial of separately formulated mRNA-1273 and mRNA-1273.529; and (7) clinically representative bivalent mRNA-1273.222 vaccine, which is a 1:1 mix in the vial of separately formulated mRNA-1273 and mRNA-1273.045. All mRNAs are formulated into a mixture of 4 lipids: SM-102, cholesterol, DSPC, and PEG2000-DMG. Preclinical mRNA vaccines were prepared with the same method as the Good Manufacturing Practice for clinical vaccines. In some experiments, clinically representative mRNA-1273.214 and mRNA-1273.222 were used.

### Viral antigens

Recombinant soluble S and RBD proteins from Wuhan-1, BA.1, and BA.5 SARS-CoV-2 strains were expressed as described^43,44^. Recombinant proteins were produced in Expi293F cells (ThermoFisher) by transfection of DNA using the ExpiFectamine 293 Transfection Kit (ThermoFisher). Supernatants were harvested 3 days post-transfection, and recombinant proteins were purified using Ni-NTA agarose (ThermoFisher), then buffer exchanged into PBS and concentrated using Amicon Ultracel centrifugal filters (EMD Millipore). SARS-CoV-2 B.1.617.2 RBD protein was purchased from Sino Biological (Cat. # 40592-V08H90).

### ELISA

Assays were performed in 96-well microtiter plates (ThermoFisher Scientific) coated with 100 µL of recombinant Wuhan-Hu-1 spike (S-2P), BA.1 spike (S-2P.529), or BA.4/BA.5 spike (S-2P.045) proteins. Plates were incubated at 4°C overnight and then blocked for 1 hour at 37°C using SuperBlock (ThermoFisher Scientific, Cat. # 37516), and then washed four times with PBS 0.05% Tween-20 (PBST). Serum samples were serially diluted in 5% bovine serum albumin in TBS (Boston BioProducts, Cat. # IBB-187), added to plates, incubated for 1 h at 37°C, and then washed four times with PBST. Goat anti-mouse IgG-HRP (Southern Biotech Cat. #1030-05) was diluted in 5% bovine serum albumin in TBS before adding to the wells and incubating for 1 h at 37°C. Plates were washed four times with PBST before the addition of TMB substrate (ThermoFisher Scientific, Cat. # 34029). Reactions were stopped by the addition of TMB stop solution (Invitrogen, Cat. # SS04). The optical density (OD) measurements were taken at 450 nm, and titers were determined using a 4-parameter logistic curve fit in Prism Version 9 (GraphPad 112 Software, Inc.) and defined as the reciprocal dilution at an OD of approximately 450 of 1 (normalized to a mouse standard on each plate).

### Focus reduction neutralization test

Serial dilutions of sera were incubated with 10^2^ focus-forming units (FFU) of WA1/2020 D614G, B.1.617.2, BA.1, or BA.5 for 1 h at 37°C. Antibody-virus complexes were added to Vero-TMPRSS2 cell monolayers in 96-well plates and incubated at 37°C for 1 h. Subsequently, cells were overlaid with 1% (w/v) methylcellulose in MEM. Plates were harvested 30 h (WA1/2020 D614G and B.1.617.2) or 70 h (BA.1 and BA.5) later by removing overlays and fixed with 4% PFA in PBS for 20 min at room temperature. Plates were washed and sequentially incubated with a pool (SARS2-02, -08, -09, -10, -11, -13, -14, -17, -20, -26, -27, -28, -31, -38, -41, -42, -44, -49,, -57, -62, -64, -65, -67, and -71^45^) of anti-S murine antibodies (including cross-reactive mAbs to SARS-CoV) and HRP-conjugated goat anti-mouse IgG (Sigma Cat # A8924, RRID: AB_258426) in PBS supplemented with 0.1% saponin and 0.1% bovine serum albumin. SARS-CoV-2-infected cell foci were visualized using TrueBlue peroxidase substrate (KPL) and quantitated on an ImmunoSpot microanalyzer (Cellular Technologies).

### VSV pseudovirus neutralization assay

Codon-optimized full-length spike genes (Wuhan-1 with D614G, BA.2.75, BA.1, and BA.5) were cloned into a pCAGGS vector. Spike genes contained the following mutations: (a) BA.2.75; T19I, Δ24-26, A27S, G142D, K147E, W152R, F157L, I210V, V213G, G257S, G339H, S371F, S373P, S375F, T376A, D405N, R408S, K417N, N440K, G446S, N460K, S477N, T478K, E484A, Q498R, N501Y, Y505H, D614G, H655Y, N679K, P681H, N764K, D796Y, Q954H, N969K (b) BA.1: A67V, Δ69-70, T95I, G142D/ΔVYY143-145, ΔN211/L212I, ins214EPE, G339D, S371L, S373P, S375F, K417N, N440K, G446S, S477N, T478K, E484A, Q493R, G496S, Q498R, N501Y, Y505H, T547K, D614G, H655Y, N679K, P681H, N764K, D796Y, N856K, Q954H, N969K, L981F; and (c) BA.4/5: T19I, Δ24-26, A27S, Δ69-70, G142D, V213G, G339D, S371F, S373P, S375F, T376A, D405N, R408S, K417N, N440K, L452R, S477N, T478K, E484A, F486V, Q498R, N501Y, Y505H, D614G, H655Y, N679K, P681H, N764K, D796Y, Q954H, N969K. To generate VSVΔG-based SARS-CoV-2 pseudovirus, BHK-21/WI-2 cells were transfected with the spike expression plasmid and infected by VSVΔG-firefly-luciferase as previously described^46^. Vero E6 cells were used as target cells for the neutralization assay and maintained in DMEM supplemented with 10% fetal bovine serum. To perform neutralization assay, mouse serum samples were heat-inactivated for 45 min at 56°C, and serial dilutions were made in DMEM supplemented with 10% FBS. The diluted serum samples or culture medium (serving as virus only control) were mixed with VSVΔG-based SARS-CoV-2 pseudovirus and incubated at 37°C for 45 min. The inoculum virus or virus-serum mix was subsequently used to infect Vero E6 cells (ATCC, CRL-1586) for 18 h at 37°C. At 18 h post infection, an equal volume of One-Glo reagent (Promega; E6120) was added to culture medium for readout using BMG PHERastar-FSX plate reader. The percentage of neutralization was calculated based on relative light units of the virus control, and subsequently analyzed using four parameter logistic curve (Prism 8,0).

### Lentivirus-based pseudovirus neutralization assay

Neutralization of SARS-CoV-2 also was measured in a single-round-of-infection assay with lentivirus-based pseudovirus assay as previously described^47^. To produce SARS-CoV-2 pseudoviruses, an expression plasmid bearing codon-optimized SARS-CoV-2 full-length spike plasmid was co-transfected into HEK293T/17 cells (ATCC#CRL-11268) cells with packaging plasmid pCMVDR8.2, luciferase reporter plasmid pHR′CMV-Luc and a TMPRSS2 plasmid. Mutant spike plasmids were produced by Genscript. Pseudoviruses were mixed with 8 serial 4-fold dilutions of sera or antibodies in triplicate and then added to monolayers of 293T-hACE2 cells in triplicate. Three days after infection, cells were lysed, luciferase was activated with the Luciferase Assay System (Promega), and RLUs were measured at 570 nm on a Spectramax L luminometer (Molecular Devices). After subtraction of background RLU (uninfected cells), % neutralization was calculated as 100 × ([virus only control]-[virus + antibody])/[virus only control]). Dose-response curves were generated with a 5-parameter nonlinear function, and titers reported as the serum dilution or antibody concentration required to achieve ID_50_ neutralization. The input dilution of serum was 1:50, thus, 20 was the lower limit of detection. Samples that did not neutralize at the limit of detection at 50% were plotted at 25, and that value was used for geometric mean calculations. Each assay included duplicates. In addition, the reported values were the geometric mean of 2 independent assays.

### Mouse experiments

#### (a) K18hACE2 transgenic mice

Seven-week-old female K18-hACE2 mice were immunized three weeks apart with 0.25 of mRNA vaccines (control or mRNA-1273) in 50 µl of PBS via intramuscular injection in the hind leg. Animals were bled 31 weeks after the second vaccine dose for immunogenicity analysis and then boosted with PBS (no vaccine) or 0.25 µg of mRNA-1273, mRNA-1273.214, or mRNA-1273.222 vaccines. Four weeks later, K18-hACE2 mice were challenged with 10^4^ FFU of BA.5 by the intranasal route. Animals were euthanized at 4 dpi, and tissues were harvested for virological analyses.

#### (b) BALB/c mice

6 to 8-week-old female BALB/c mice were immunized three weeks apart with 1 µg of mRNA vaccines (mRNA-1273, mRNA-1273.529, mRNA-1273.045, mRNA-1273.214, or mRNA-1273.222) or PBS (in 50 µL) via intramuscular injection in the quadriceps muscle of the hind leg under isoflurane anesthesia. Blood was sampled three weeks after the first immunization and two weeks after the second immunization, and anti-spike and neutralizing antibody levels were measured by ELISA, and VSV-based or lentivirus-based pseudovirus neutralization assays.

### Measurement of viral burden

Tissues were weighed and homogenized with zirconia beads in a MagNA Lyser instrument (Roche Life Science) in 1 ml of DMEM medium supplemented with 2% heat-inactivated FBS. Tissue homogenates were clarified by centrifugation at 10,000 rpm for 5 min and stored at −80°C.

#### Viral RNA measurement

RNA was extracted using the MagMax mirVana Total RNA isolation kit (Thermo Fisher Scientific) on the Kingfisher Flex extraction robot (Thermo Fisher Scientific). RNA was reverse transcribed and amplified using the TaqMan RNA-to-CT 1-Step Kit (Thermo Fisher Scientific). Reverse transcription was carried out at 48°C for 15 min followed by 2 min at 95°C. Amplification was accomplished over 50 cycles as follows: 95°C for 15 sec and 60°C for 1 min. Copies of SARS-CoV-2 *N* gene RNA in samples were determined using a published assay^48^.

#### Viral plaque assay

Vero-TMPRSS2-hACE2 cells were seeded at a density of 1×10^5^ cells per well in 24-well tissue culture plates. The following day, medium was removed and replaced with 200 µL of clarified lung homogenate that was diluted serially in DMEM supplemented with 2% FBS. One hour later, 1 mL of methylcellulose overlay was added. Plates were incubated for 96 h, then fixed with 4% paraformaldehyde (final concentration) in PBS for 20 min. Plates were stained with 0.05% (w/v) crystal violet in 20% methanol and washed twice with distilled, deionized water.

### Cytokine and chemokine protein measurements

Lung homogenates were incubated with Triton-X-100 (1% final concentration) for 1 h at room temperature to inactivate SARS-CoV-2. Homogenates were analyzed for cytokines and chemokines by Eve Technologies Corporation (Calgary, AB, Canada) using their Mouse Cytokine Array/Chemokine Array 31-Plex (MD31) platform.

### Materials availability

All requests for resources and reagents should be directed to the corresponding authors. This includes viruses, vaccines, and primer-probe sets. All reagents will be made available on request after completion of a Materials Transfer Agreement (MTA). All mRNA vaccines can be obtained under an MTA with Moderna (contact: Darin Edwards).

### Data and code availability

All data supporting the findings of this study are available within the paper and are available from the corresponding author upon request. Any additional information required to reanalyze the data reported in this paper is available from the lead contact upon request.

### Code availability

No code was used in the course of the data acquisition or analysis.

### Statistical analysis

Significance was assigned when *P* values were < 0.05 using GraphPad Prism version 9.3. Tests, number of animals, median values, and statistical comparison groups are indicated in the Figure legends. Changes in infectious virus titer, viral RNA levels, or serum antibody responses were compared to unvaccinated or mRNA-control immunized animals and were analyzed by Kruskal-Wallis or one-way ANOVA with multiple comparisons tests, or Wilcoxon signed-rank test depending on the type of results, number of comparisons, and distribution of the data.

