## Supplemental Table S1 for "Bivalent SARS-CoV-2 mRNA vaccines increase breadth of neutralization and protect against the BA.5 Omicron variant"

**Supplementary Table S1. Cytokine and chemokine concentrations in the lungs of immunized K18-hACE2 mice challenged with BA.5**

|  | **Concentration (mean ± SD (pg/mL))** | | | | | |
| --- | --- | --- | --- | --- | --- | --- |
|  | **Naive** | **CTRL-mRNA**  **+CTRL-mRNA** | **mRNA-1273**  **+ PBS** | **mRNA-1273**  **+ mRNA-1273** | **mRNA-1273**  **+ mRNA-1273.214** | **mRNA-1273**  **+ mRNA-1273.222** |
| **G-CSF** | 0.67 + 0 | 4.51 + 3.94 | 6.05 + 3.39 | 1.67 + 1.7 | 0.7 + 0.09 | 0.88 + 0.41 |
| **GM-CSF** | 0.68 + 0 | 4.47 + 4.42 | 6.19 + 4.02 | 2.17 + 3.15 | 0.68 + 0 | 0.68 + 0 |
| **IFN-γ** | 0.86 + 0.25 | 2.1 + 0.87 | 5.29 + 1.64 | 1.95 + 1.22 | 0.89 + 0.4 | 1.09 + 0.33 |
| **IL-1α** | 26.01 + 10 | 40.1 + 5.53 | 41.5 + 6.74 | 37.42 + 10.96 | 40.95 + 13.33 | 31.28 + 8.51 |
| **IL-1β** | 0.63 + 0 | 1.11 + 0.47 | 1.71 + 0.73 | 0.87 + 0.47 | 0.64 + 0.02 | 0.66 + 0.09 |
| **IL-2** | 2.24 + 1.05 | 3.41 + 1.24 | 7.15 + 1.69 | 5.31 + 1.78 | 5.76 + 2.53 | 4.18 + 1.1 |
| **IL-6** | 0.66 + 0 | 55.52 + 30.11 | 93.72 + 75.69 | 26.24 + 54.25 | 1.34 + 1.13 | 1.17 + 0.78 |
| **IL-7** | 1.03 + 0.26 | 0.94 + 0.19 | 1.32 + 0.61 | 1.17 + 0.85 | 0.88 + 0.33 | 0.73 + 0.13 |
| **IL-9** | 32.58 + 10.22 | 69.88 + 22.19 | 79.83 + 15.86 | 61.76 + 20.09 | 33.6 + 19.44 | 61.18 + 11.29 |
| **IL-15** | 0.64 + 0 | 1.88 + 0.95 | 3.26 + 1.43 | 1.01 + 0.6 | 0.72 + 0.16 | 0.67 + 0.09 |
| **IP-10 (CXCL10)** | 20.43 + 7.69 | 818.41 + 441.1 | 701.38 + 371.91 | 142.97 + 185.73 | 25.82 + 25.14 | 21.57 + 8.55 |
| **KC (CXCL1)** | 18.57 + 6.49 | 66.27 + 29.7 | 163.22 + 101.41 | 57.38 + 68.75 | 36.36 + 22.06 | 42.2 + 27.88 |
| **LIF** | 0.65 + 0 | 3.52 + 1.86 | 5.15 + 2.34 | 2.13 + 2.71 | 0.65 + 0 | 0.66 + 0.03 |
| **MCP-1 (CCL2)** | 12.14 + 6.49 | 434.73 + 186.97 | 581.57 + 367.7 | 125.52 + 167.45 | 30.2 + 20.26 | 36.04 + 27.97 |
| **M-CSF** | 3.27 + 0.36 | 3.82 + 1.12 | 6.02 + 2.15 | 4.04 + 3.59 | 1.98 + 0.55 | 2.69 + 1.01 |
| **MIG (CXCL9)** | 29.05 + 14.15 | 189.05 + 99.95 | 283.48 + 106.89 | 112.11 + 150.27 | 11.38 + 7.35 | 21.11 + 12.97 |
| **MIP-1α (CCL3)** | 11.19 + 5.82 | 72.68 + 20.96 | 110.78 + 45.71 | 111.08 + 103.06 | 126.83 + 125.77 | 26.81 + 11.92 |
| **MIP-1β (CCL4)** | 9.9 + 5.04 | 67.18 + 16.31 | 67.62 + 35.94 | 22.28 + 28.9 | 5.4 + 3.62 | 6.66 + 5.9 |
| **MIP-2 (CXCL2)** | 23.02 + 8.4 | 38.02 + 14.92 | 45.92 + 9.3 | 37.98 + 14.58 | 26.7 + 8.97 | 35.34 + 12.64 |
| **TNF-α** | 0.59 + 0 | 1.65 + 0.75 | 2.34 + 1.03 | 0.96 + 0.71 | 0.71 + 0.27 | 0.59 + 0 |

Seven-week-old female K18-hACE2 mice were immunized with 0.25 μg of control mRNA or mRNA-1273, boosted 31 weeks later with PBS, 0.25 μg of control mRNA, or 0.25 μg of clinically representative versions of mRNA-1273, mRNA-1273.214, or mRNA-1273.222 vaccines, and then one month later challenged via intranasal route with 10^4^ focus-forming units (FFU) of BA.5. A separate set of naïve K18-hACE2 mice were used for comparison. Cytokine and chemokine levels in lung homogenates are expressed as mean ± standard deviation in pg/mL (2 experiments, n = 8-10 per group).
