## Supplementary figures and images for "Bivalent SARS-CoV-2 mRNA vaccines increase breadth of neutralization and protect against the BA.5 Omicron variant"

### Extended Data Figure 1

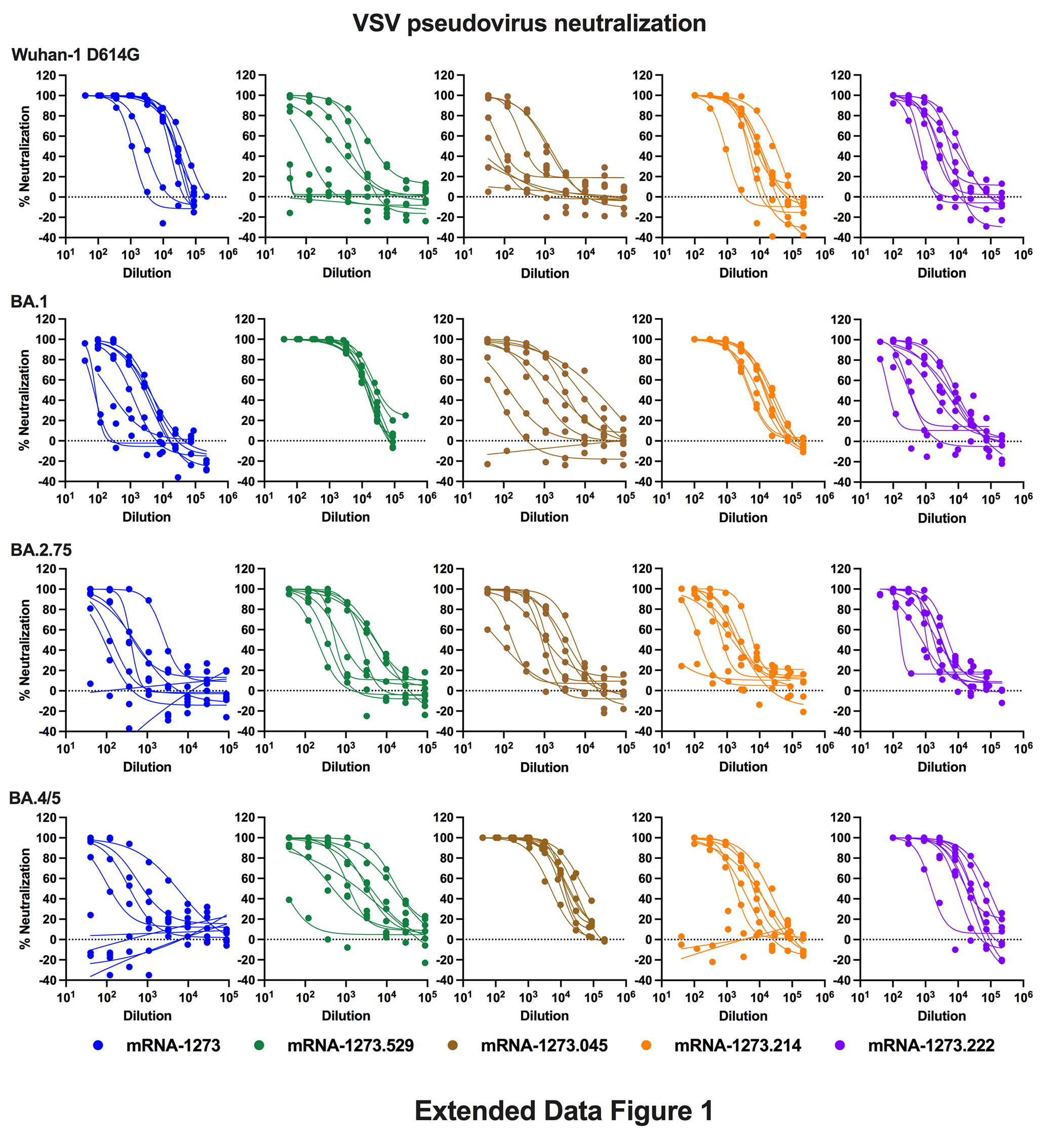

### Extended Data Figure 2

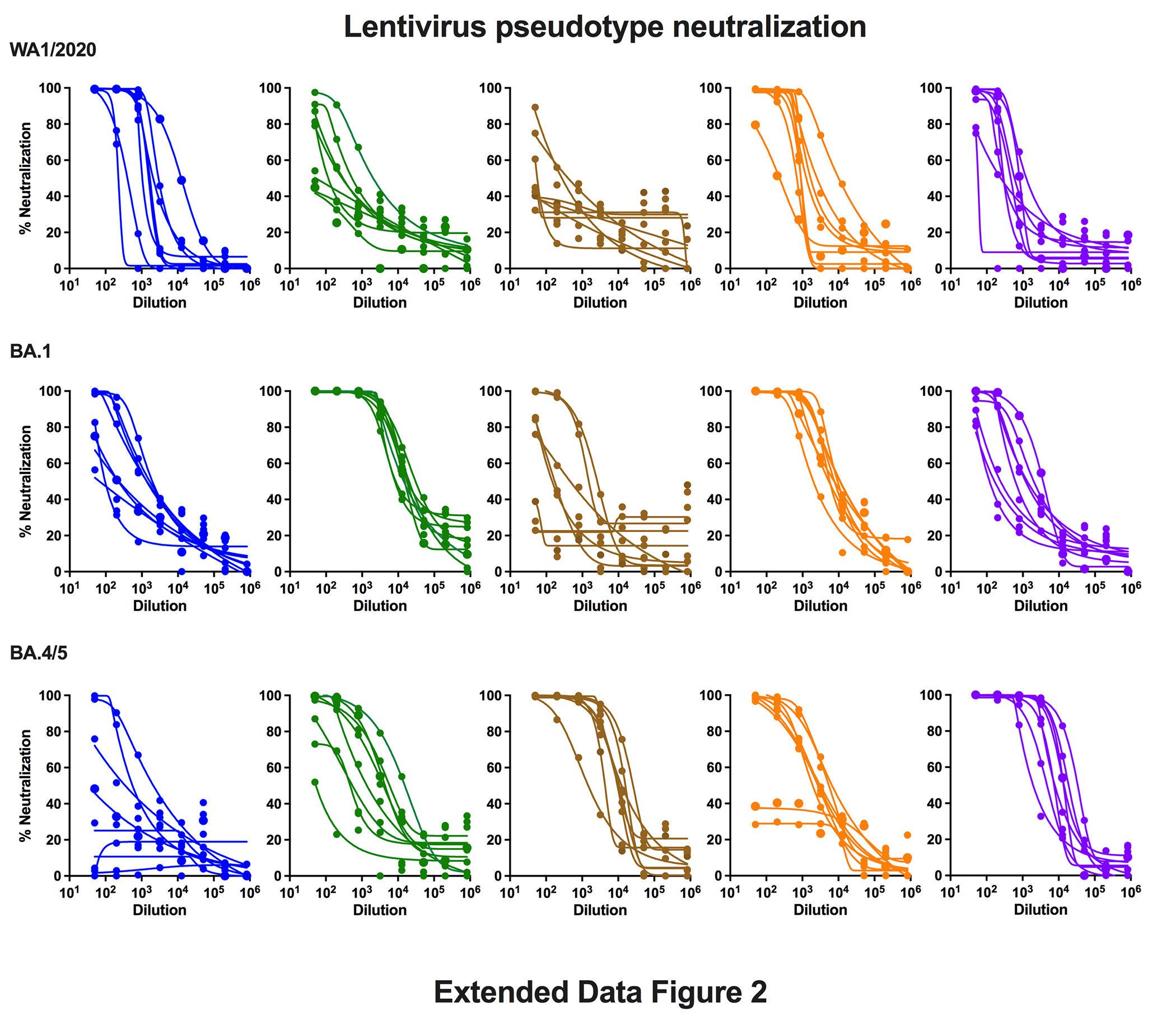

### Extended Data Figure 3

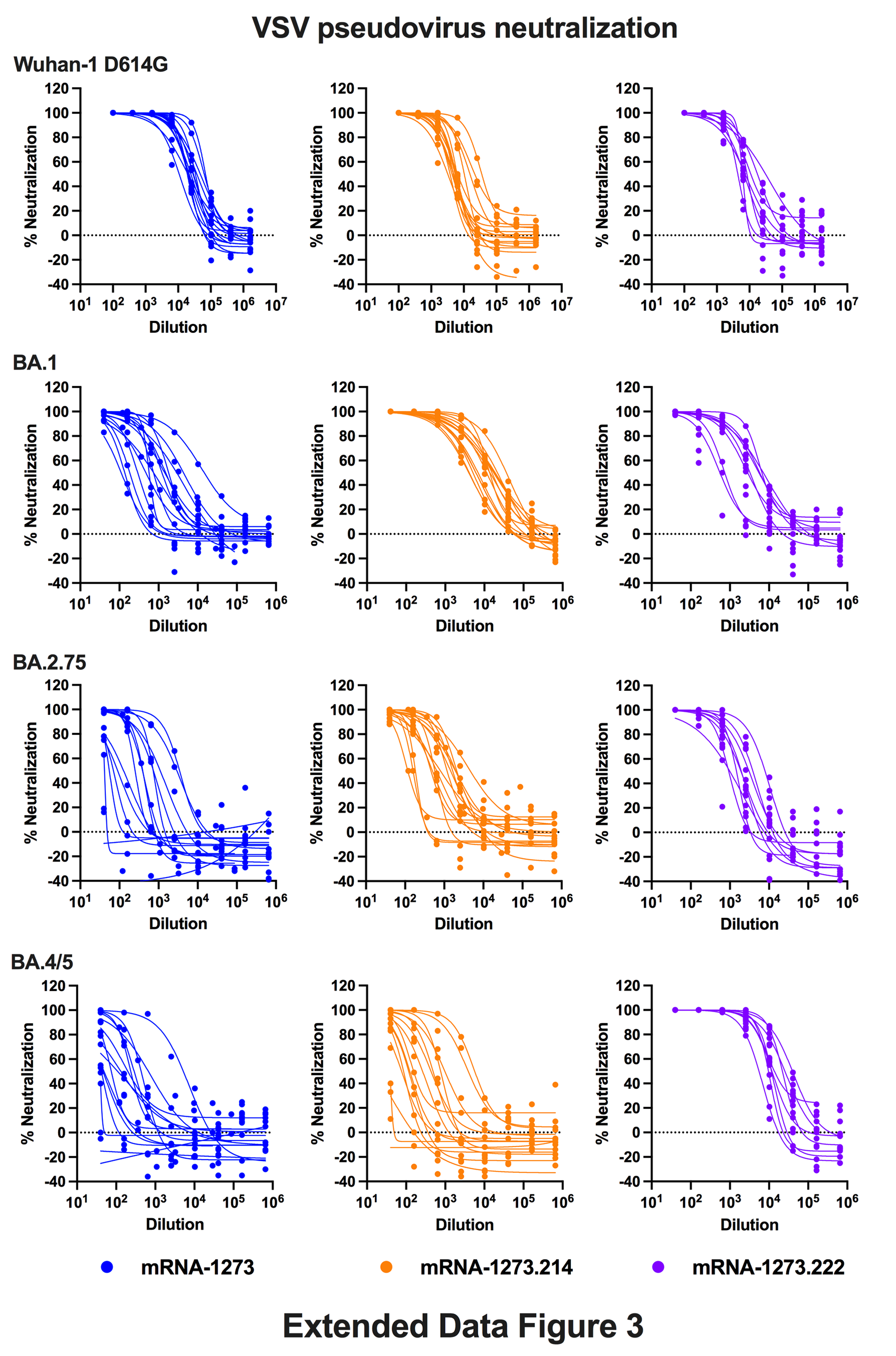

### Extended Data Figure 4

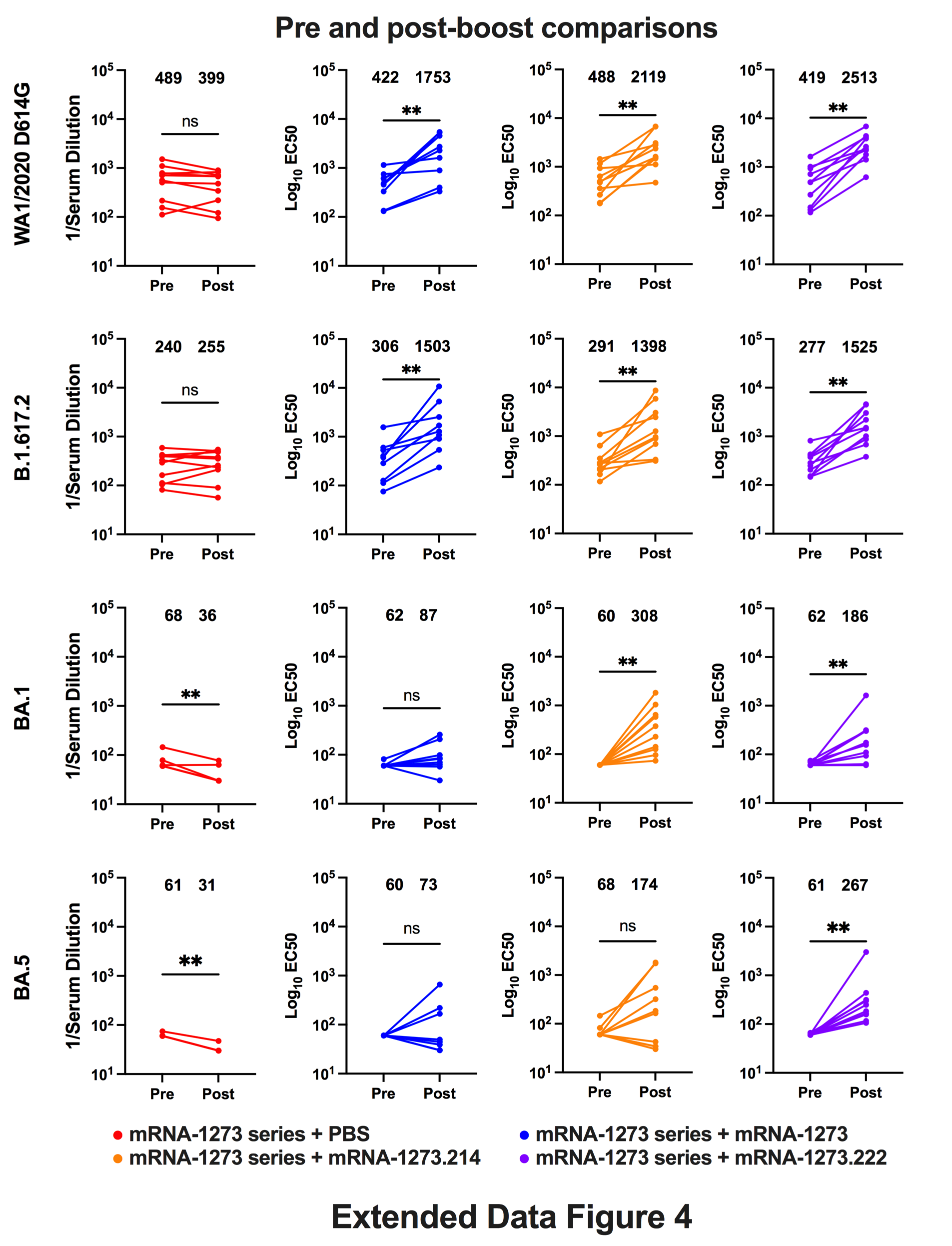

### Extended Data Figure 5

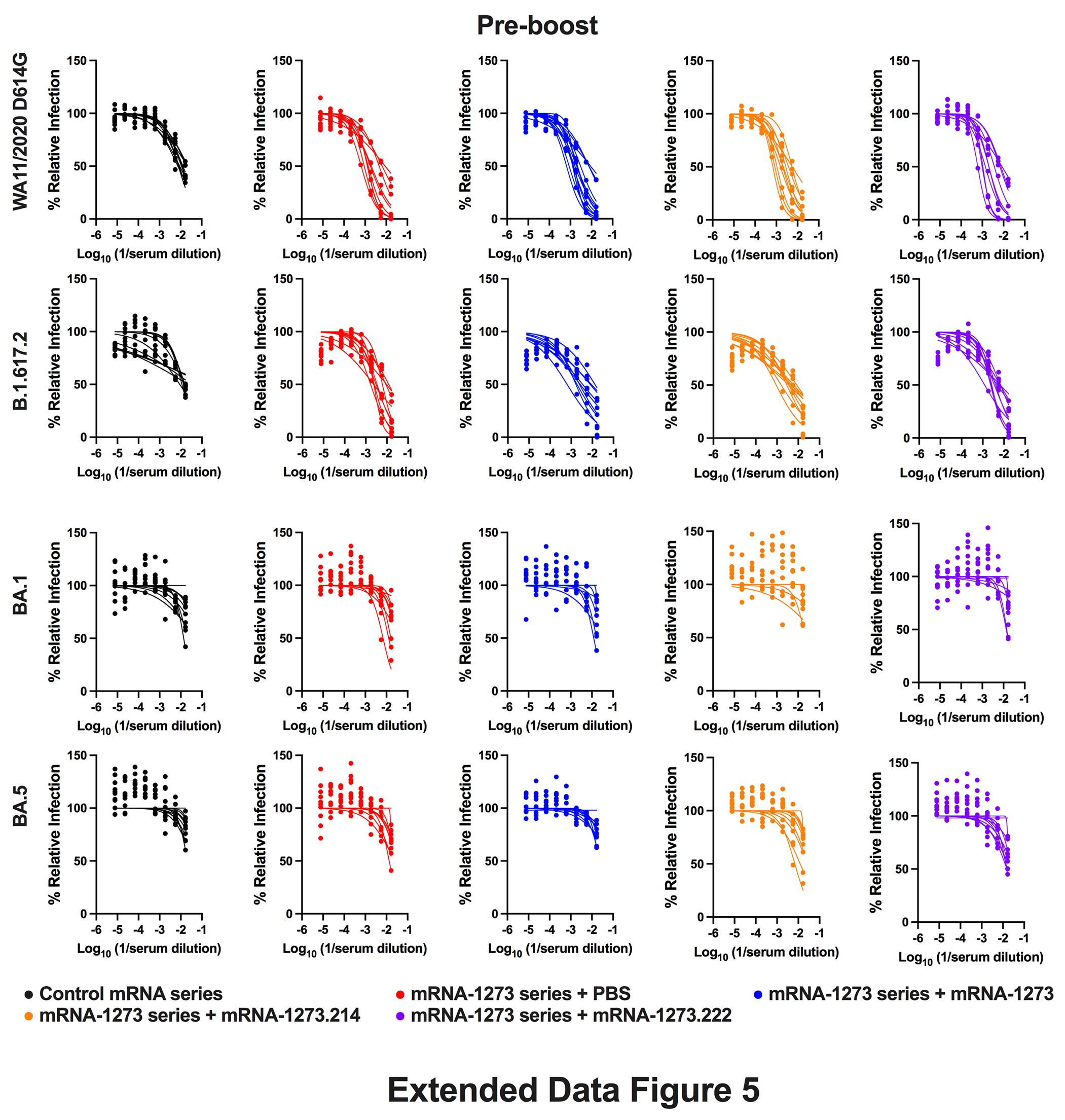

### Extended Data Figure 6

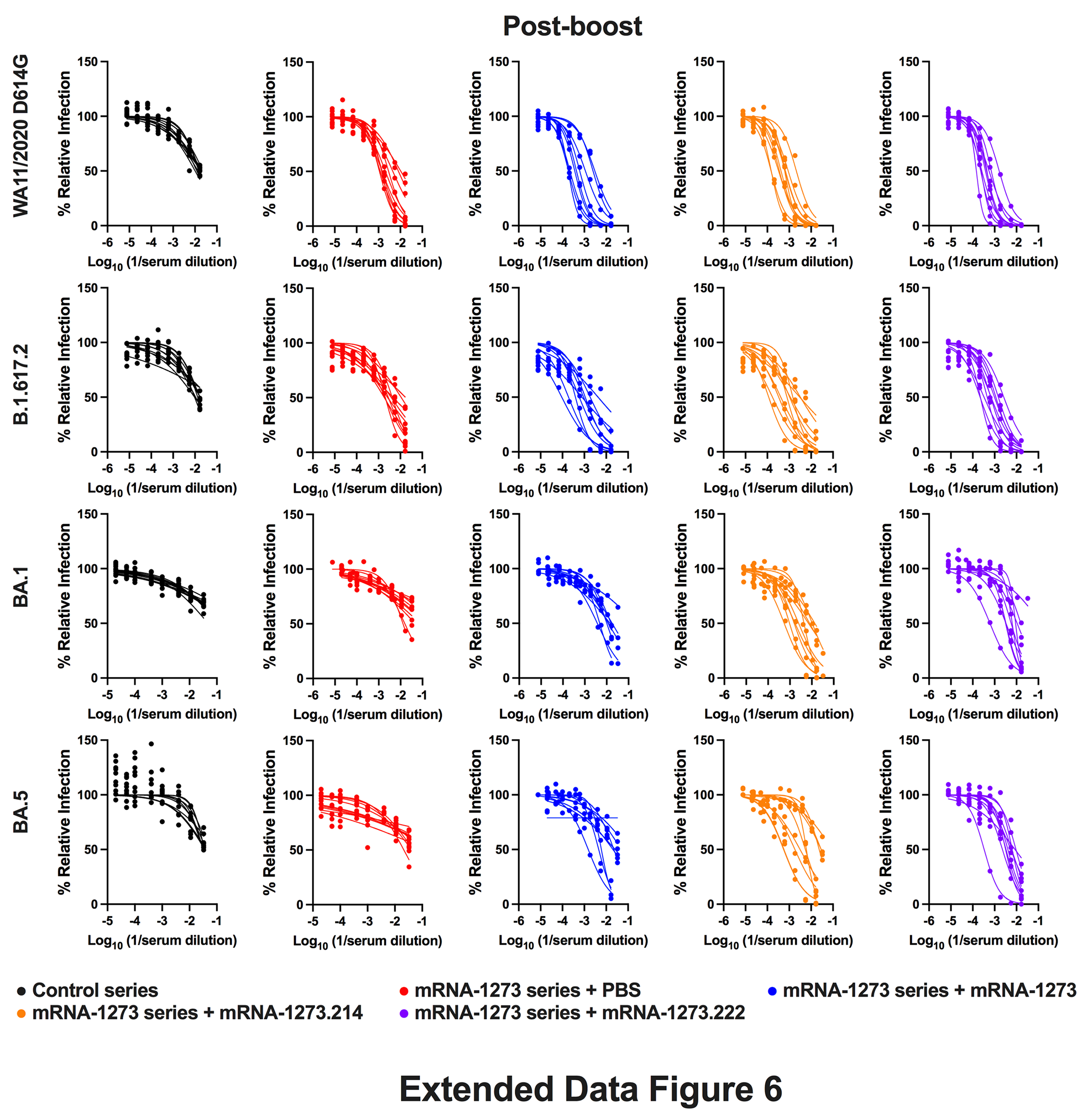
